# Structure of Transmembrane AMPA Receptor Regulatory Protein Subunit γ2

**DOI:** 10.1101/2023.11.28.569079

**Authors:** W. Dylan Hale, Alejandra Montaño Romero, Richard L. Huganir, Edward C. Twomey

## Abstract

Transmembrane AMPA receptor regulatory proteins (TARPs) are claudin-like proteins that tightly regulate AMPA receptors (AMPARs) and are fundamental for excitatory neurotransmission. We used cryo-electron microscopy (cryo-EM) to reconstruct the 36 kDa TARP subunit γ2 to 2.3 Å and reveal the structural diversity of TARPs. Our data reveals critical motifs that distinguish TARPs from claudins and define how sequence variations within TARPs differentiate subfamilies and their regulation of AMPARs.

---

Information transfer in the brain occurs at specialized cellular junctions known as synapses, which act as neuronal communication hubs^1^. Most synapses are glutamatergic, where a pre-synaptic neuron releases glutamate (Glu), and a post-synaptic neuron receives Glu. AMPARs in the post-synaptic membrane bind Glu and initiate depolarization of the post-synaptic neuron through their Glu-gated cation channels^1,2^. TARPs are auxiliary subunits that regulate the trafficking, gating kinetics, and pharmacology of AMPARs^2,3^.

TARP regulatory subunits tightly regulate AMPAR function in the post-synaptic membrane, which is a critical aspect of the brain’s ability to fine tune information processing^1–3^. There are six TARP subtypes (TARPγ2, γ3, γ4, γ5, γ7, γ8), split into type-I (TARPγ2, γ3, γ4, γ8) and type-II (TARPγ5, γ7) families. Generally, TARPs increase the conductance of AMPARs, but type-I TARPs slow desensitization and deactivation kinetics, while type-II TARPs appear to have a negative effect on gating when compared to type-I TARPs^2^. Furthermore, structural differences between TARPs in the same class underlie sensitivity to certain classes of drugs targeted to AMPAR-TARP complexes. Since the first TARP was identified a quarter century ago (TARPγ2, also known as stargazin)^4^, TARPs have been recognized as an indispensable component of synaptic function^1,2^. Yet, the structural details of how TARPs regulate AMPARs remain ambiguous.

Cryo-EM studies of TARP subunits have advanced our understanding of TARP structure in the context of AMPAR complexes, but intermediate resolution has historically precluded *de novo* building of TARP structures^5–14^. X-ray crystallography structures of TARP homologs, such as claudins, have been indispensable for modeling TARPs^15^. Claudins are cellular junction proteins that form paracellular barriers between epithelial and endothelial cells and are functionally distinct from TARPs^16^. The reliance on claudin structures for TARP modeling has hampered identification of distinct structural features that 1) differentiate TARPs from claudins and 2) explain the regulatory potential of TARPs for AMPARs. Here, we use cryo-EM to determine the structure of the prototypical TARP, TARPγ2. We identify new motifs in TARPγ2 that distinguish TARP classes from one another and further differentiate TARPs from Claudins. These structural features likely underlie modulatory effects exhibited by TARPs on AMPAR gating.

We reconstructed the 3D architecture of TARPγ2 to an overall resolution of 2.3 Å (2.0 Å – 2.5 Å locally; **Extended Data Fig. 1**). Our data enables us to build most of the transmembrane domain (TMD) and extracellular domain (ECD) *de novo* (**Fig. 1a**). The high resolution of our reconstruction enables identification of multiple distinct structural features in the TARPγ2 extracellular domain (ECD), which sits atop its tetraspanin transmembrane (TM) helical bundle comprised of transmembrane (TM) helices TM1-4 (**Fig. 1a**). The ECD is comprised of a five-stranded β-sheet and a single extracellular helix (ECH) that immediately precedes TM2. A previously identified disulfide bridge (DSB) between β3 (C67) and β4 (C77) strands in the ECD stabilizes the TARPγ2 ECD (**Fig. 1b**) and is conserved across all TARPs and the TARP-like claudins.

**Figure 1.**
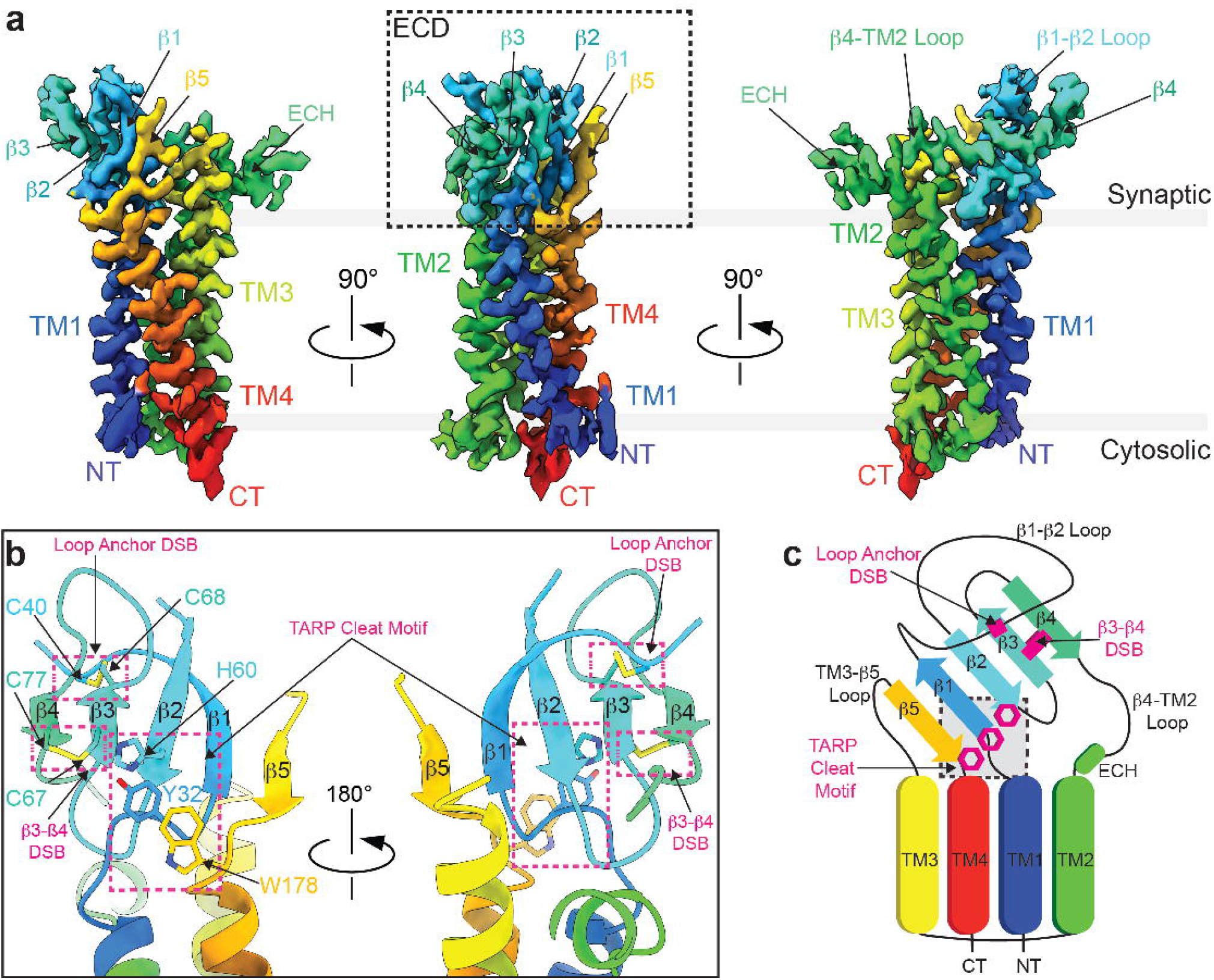
Structure of TARPγ2. a) Cryo-EM map of TARPγ2, colored rainbow from N-terminus, NT (blue) to C-terminus, CT (red). b) Extracellular portion of the TARPγ2 model showing the β3-β4 DSB, loop anchor DSB, and TARP cleat. c) Cartoon schematic of TARPγ2 structure highlighting key structural features that rigidify the entire ECD atop the tetraspanin TMD, colored as in panel a.

What makes TARPγ2, and all TARPs unique from claudins? We identify two new moieties in our reconstruction of TARPγ2 that distinguish TARPs from claudins. First, a π-π-π stack secures the TARPγ2 ECD atop the TARPγ2 TMD (**Fig. 1b**). This is formed by H60 (from β2), Y32 (TM1-β1 loop), and W178 (TM4). We term this the TARP cleat motif because it helps to fasten the ECD to the TMD. We also identified a second DSB in the ECD. This DSB, the loop anchor DSB, anchors the β1-β2 loop onto the β-sheet (**Fig. 1b**). The loop anchor DSB is made between C40 in the β1-β2 loop and C68 on β3. All together, these motifs rigidify the structure of TARPγ2 by providing additional structural interactions within the ECD and between the ECD and TMD (**Fig. 1c**).

How conserved are these motifs? The TARP cleat motif is conserved in all TARPs and the TARP-like subunit germline specific gene 1-like (GSG1L) (**Fig. 2a**) but absent from all claudins (**Extended Data Fig. 2**). We also tested for conservation of the cleat motif through AlphaFold2^17^ structure prediction. This suggests that the TARP cleat motif is present in all mammalian TARPs (**Extended Data Fig. 3a**). Interestingly, while the TARP cleat motif is conserved in all TARPs, the loop anchor DSB is not (**Fig. 2a**). Structure prediction in AlphaFold2 (**Extended Data Fig. 3b**) also points to the loop anchor DSB being conserved in type-I TARPs but not in type-II TARPs. Thus, while our structure pointed us to look at the conservation of the cleat motif and loop anchor DSB, this was already predicted by AlphaFold2 (**Extended Data Fig. 3c**).

**Figure 2.**
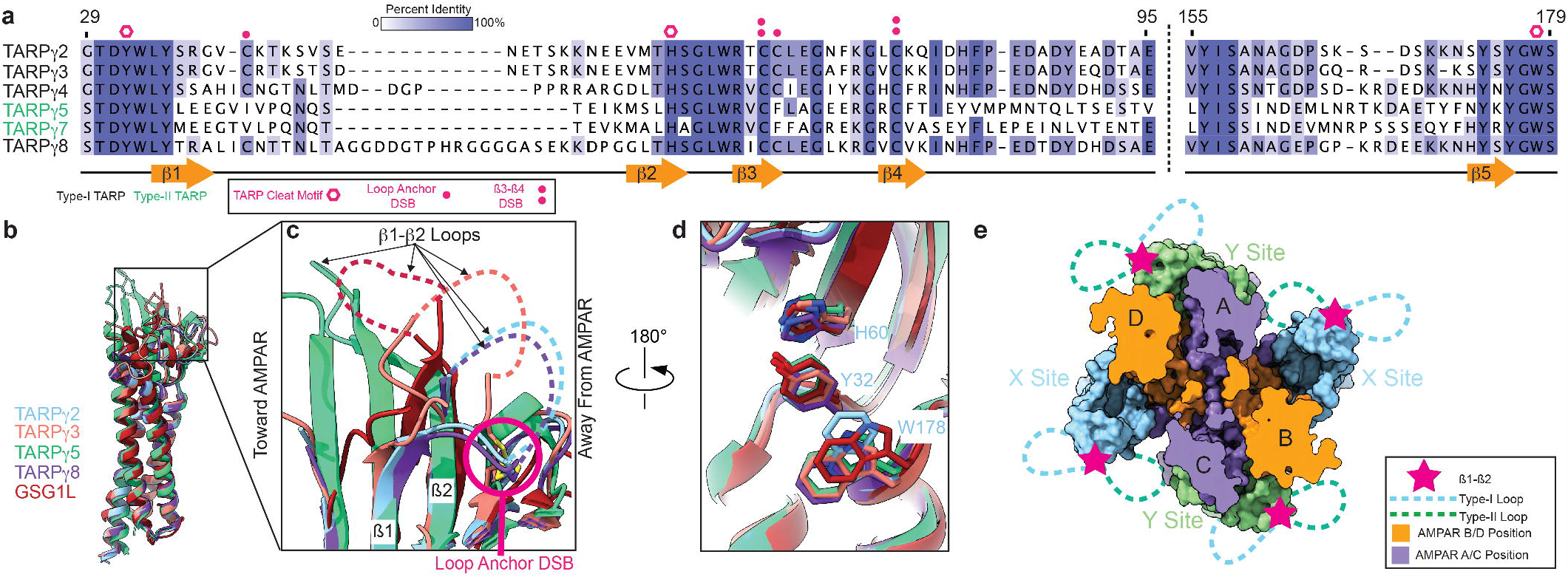
Conservation of structural features among TARP family members. a) Multiple sequence alignment demonstrating the relative conservation of the TARP Cleat Motif, β3-β4 DSB, and Loop Anchor DSB between TARP family members. Loop Anchor DSB is unique to type-I and excluded from type-II TARPs. b) Alignment of TARPγ2 structure with other TARP family members (TARPγ3, PDB: 8C2H; TARPγ5, PDB: 7RZ5; TARPγ8, PDB: 8AYN; GSG1L, PDB: 7RZ9). c) Zoomed in view of TARP extracellular domains illustrating differing orientations in the β1-β2 loops. d) View of the TARP cleat motif illustrating conservation among all TARP family members. e) Model of predicted β1-β2 loop orientations between type-I and type-II TARPs illustrating distinct potential contacts between TARP subtypes and AMPARs.

Surprisingly, the TARP cleat motif and loop anchor DSB are within previous TARP structures but not identified. Previously determined structures of TARPs are overall like our structure of TARPγ2 (**Fig. 2b**), and the loop anchor DSB is within structures of TARPγ3^18^ and TARPγ8^11,12,19^, and even previously published structures of TARPγ2^6^. However, it is absent, as expected, in the structure of the type-II TARP, TARPγ5^20,21^ (**Fig. 2c**) and the TARP-like subunit GSG1L^7,20^ (**Fig. 2c**). In contrast, the TARP cleat motif is conserved in all TARPγ3, γ5, and γ8 subunit structures as well as GSG1L^11,18,20^ (**Fig. 2d**). Thus, we suggest expanding the type-II family of TARPs to include the GSG1L subunit. We hypothesize that these structural details and their conservation were previously missed because of a lack of structural resolution.

The dichotomy in β1-β2 loop organization between type-I and type-II TARPs has significant functional implications. For example, type-II TARPs lack the loop anchor DSB and have been observed to directly interact with AMPAR subunits that are in the A and C positions when they occupy the “X” auxiliary subunit site^7,20^ (**Fig. 2e**). However, we expect that this is not possible for type-I TARPs in the X site given the presence of the loop anchor DSB, which locks in the β1-β2 loop in an orientation away from the A and C AMPAR subunit positions. However, if a type-I TARP occupies the “Y” TARP position (**Fig. 2e**), modulation of the AMPAR at subunit positions B or D by the β1-β2 loop is likely possible despite the loop anchor DSB, and is supported by observations in cryo-EM studies of type-I TARPs in complex with AMPARs^18^. Given the extreme conformational changes associated with AMPAR gating, the stark difference in the presence or absence of the loop anchor DSB within type-I TARPs versus type-II TARPs potentially explains differences in electrophysiology experiments between chimeric constructs of the β1-β2 loop in type-I and type-II TARPs.

The TARP cleat motif plays a significant role in distinguishing TARPs from claudins. Both TARPs and claudins share the same overall structural fold (i.e., tetraspanin with a five-stranded extracellular β-sheet). However, claudins have strong oligomerization properties, where they self-oligomerize to form paracellular barriers. A similar phenomenon has not been reported for TARP proteins. We hypothesize that the TARP cleat motif plays a role in preventing oligomerization in TARPs, enabling their complexation with AMPARs and other synaptic proteins.

In sum, we report the structure of TARPγ2, and how the newly identified structural features may account for critical functional differences between TARPs that tune AMPAR function throughout the central nervous system. In addition, we precisely define how TARPs are differentiated from claudins, which may explain the critical point of divergence between the structurally related proteins that are functionally distinct. Our findings provide a new framework for future studies to understand the function of TARPs and new foundations to target TARPs in structure-based drug design against AMPAR-related neurological disorders.

## Methods

### Construct design, protein expression, and purification

Mouse TARPγ2 was covalently fused to the rat AMPAR subunit GluA2, expressed, and purified as described in the preprint Hale, *et al. Biorxiv* 2023 (BIORXIV/2023/569057).

### Cryo-EM Sample Preparation and Data Collection

Cryo-EM samples were prepared and collected as described in the preprint Hale, *et al. Biorxiv* 2023 (BIORXIV/2023/569057).

### Image Processing

The initial stages of cryo-EM sample preparation were carried out as in the preprint Hale, *et al. Biorxiv* 2023 (BIORXIV/2023/569057). After generation of a 2.80 Å AMPAR-TARPγ2 local map (Extended Data Fig. 1a), symmetry expansion was used to refine the structure of TARPγ2. To achieve this, we applied C4 symmetry to the AMPAR-TARP particles (Extended Data Fig. 1a). We masked one TARPγ2 in the AMPAR-TARPγ2, then inverted this mask, and subtracted the inverted mask from all particle images. We then used the subtracted particle images, coupled with the original TARPγ2 mask (non-inverted) applied to the complete AMPAR-TARPγ2 complex cryo-EM map reference to refine the final cryo-EM reconstruction of TARPγ2 (Extended Data Fig. 1b).

### Model building, refinement, and structural analysis

Coot^22^ was used to build a polyalanine chain into TARPγ2 map. Bulky resides from sequence information were used to anchor the building. A previously determined structure of TARPγ2 (pdb 5WEO) and a structure predicted from AlphaFold2 (AlphaFold Protein Structure Database, #AF-O88602) were used as reference. Isolde^23^ and Phenix^24^ were used to refine the model. Quality of the model was assessed with MolProbity^25^. Visualizations and domain measurements were performed in ChimeraX^26^. Software was compiled and accessed via the SBGrid Consortium^27^.

### Sequence Analysis

All sequence alignments were done with ClustalW^28^ and analyzed in Jalview^29^.

### Structure Prediction

TARP structure predictions of TARPγ2, γ3, γ4, γ5, γ7, γ8 of human, rat, mouse species were used from AlphaFold2^17^. For each TARP subunit structure prediction, the respective amino acids corresponding to the cleat motif and disulfide bridge were determined. Cleat motif measurements were taken by calculating the distance between the Cα’s of histidine to tyrosine and Cα’s of tyrosine to tryptophan. Calculations were performed using the Biopython.PDB package.

AlphaFold2 accession numbers of models: AF-Q9Y698, AF-A0JNG9, AF-O88602, AF-Q71RJ2, AF-Q9JJV5, AF-Q0VD05, AF-O60359, AF-Q8VHX0, AF-A0A3Q1LKG2, AF-Q9JJV4, AF-Q8VHW9, AF-Q9UBN1, AF-E1BEI3, AF-Q8VHW4, AF-Q8VHW8, AF-Q9UF02, AF-E1BIG3, AF-P62956, AF-P62957, AF-P62955, AF-Q8WXS5, AF-F1MV40, AF-Q8VHW2, AF-Q8VHW5.

## Data Availability

All cryo-EM reconstructions will be deposited into the Electron Microscopy Data Bank (EMDB) upon publication. All micrographs from the IS-1 and IS-2 datasets will be deposited into the Electron Microscopy Public Image Archive (EMPIAR) upon publication. All structural models generated from cryo-EM will be deposited in the Protein Data Bank upon publication.

## Acknowledgements

We thank members of the Twomey and Huganir labs for insightful discussions. All cryo-EM data was collected at the Beckman Center for Cryo-EM at Johns Hopkins with assistance from D. Sousa and D. Ding.

## Author Contributions

E.C.T. and R.L.H. supervised all aspects and planning of this research. E.C.T., A.M.R., and W.D.H. designed the project. E.C.T. and W.D.H. wrote the manuscript with input from all authors. W.D.H. prepared samples for cryo-EM, collected cryo-EM data, processed cryo-EM data, analyzed data, and built models with E.C.T. A.M.R. assisted with structural analysis, structure prediction, model building, data analysis, structural analysis, and in uncovering the conserved TARP motifs.

## Cryo-EM data collection, refinement and validation statistics

|  | TARP<sub>γ</sub>2<br>(EMDB-xxxx)<br>(PDB xxxx) |
| --- | --- |
| **Data collection and processing** |  |
| Magnification | 130,000x |
| Voltage (kV) | 300 |
| Electron exposure (e<sup>-</sup>/Å<sup>2</sup>) | 40 |
| Defocus range (μm) | -1.0 – 2.6 |
| Pixel size (Å) | 0.93 |
| Symmetry imposed | C1 |
| Initial particle images (no.) | 123,729 |
| Final particle images (no.) | 494,916 |
| Map resolution (Å) | 2.32 |
| FSC = 0.143 |  |
| Map resolution range (Å) | 2 – 4 |
| **Refinement** |  |
| Initial model used (PDB code) | N/A |
| Model resolution (Å) | 2.3 |
| FSC = 0.143 |  |
| Model resolution range (Å) | 2.1 – 3.7 |
| Map sharpening *B* factor (Å<sup>2</sup>) | -54.8 |
| Model composition |  |
| Non-hydrogen atoms | 2678 |
| Protein residues | 172 |
| Ligands | 0 |
| *B* factors (Å<sup>2</sup>) |  |
| Protein | 0.00/27.91/5.8 |
| Ligand | N/A |
| R.m.s. deviations |  |
| Bond lengths (Å) | 0.013 |
| Bond angles (°) | 1.726 |
| Validation |  |
| MolProbity score | 0.71 |
| Clashscore | 0 |
| Poor rotamers (%) | 1.44 |
| Ramachandran plot |  |
| Favored (%) | 97.56 |
| Allowed (%) | 1.83 |
| Disallowed (%) | 0.61 |

## Figure Legends

**Extended Data Figure 1.**
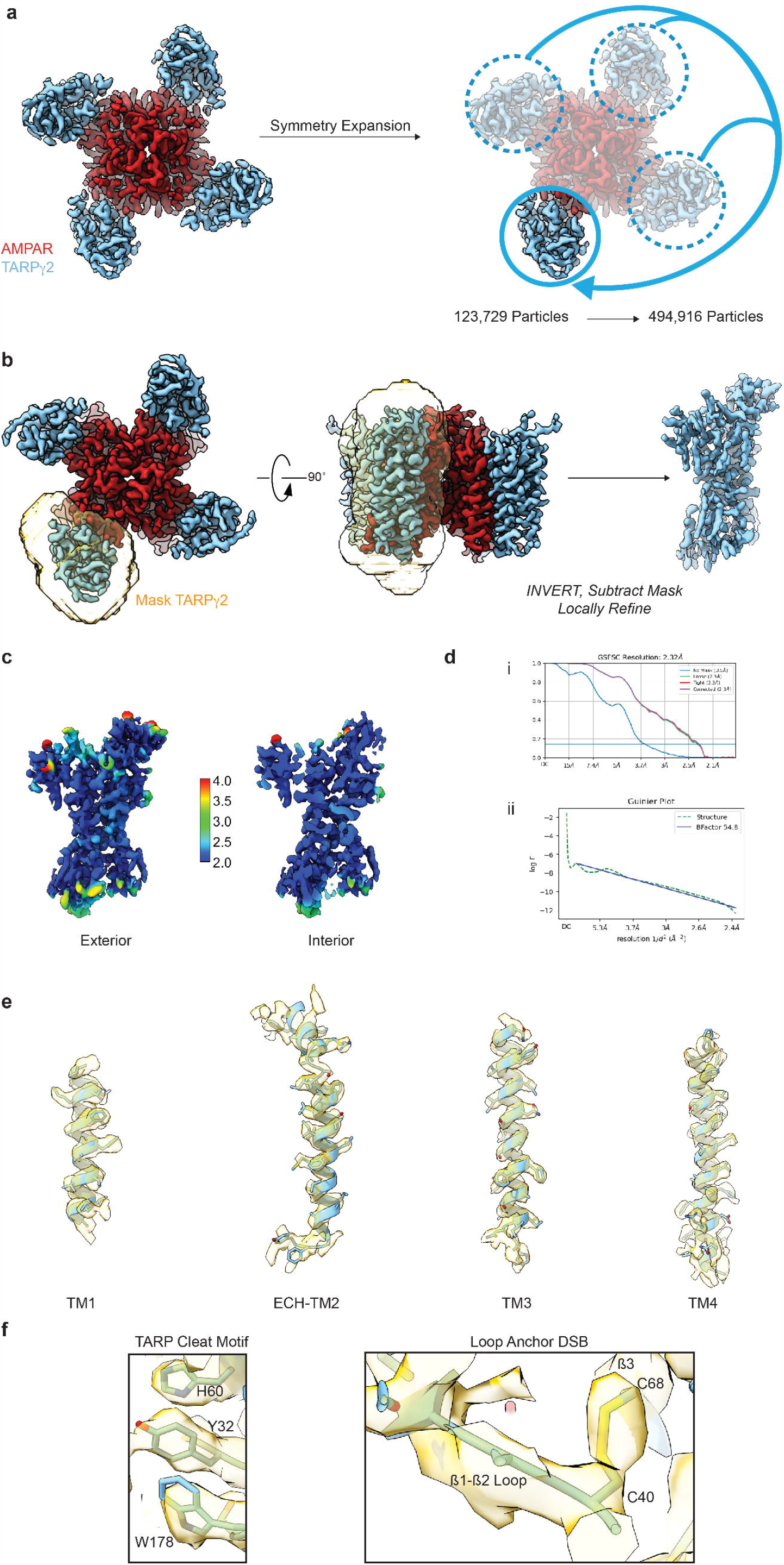
Details of TARPγ2 data processing workflow. a) Symmetry expansion of the GluA2-TARPγ2 assembly (from Hale et al., 2023, *BioRxiv*). b) Masking scheme for isolating symmetry-expanded TARPγ2. c) TARPγ2 cryo-EM map colored by local resolution *right:* surface of TARPγ2 reconstruction, *left:* cutaway showing resolution inside the map. d) Gold Standard Fourier Shell Correlation and Guinier Plots for TARPγ2. e) Model fit to cryo-EM map of the four TARPγ2 TM helices. f) Cryo-EM map around the TARP Cleat motif and the Loop Anchor DSB.

**Extended Data Figure 2.**
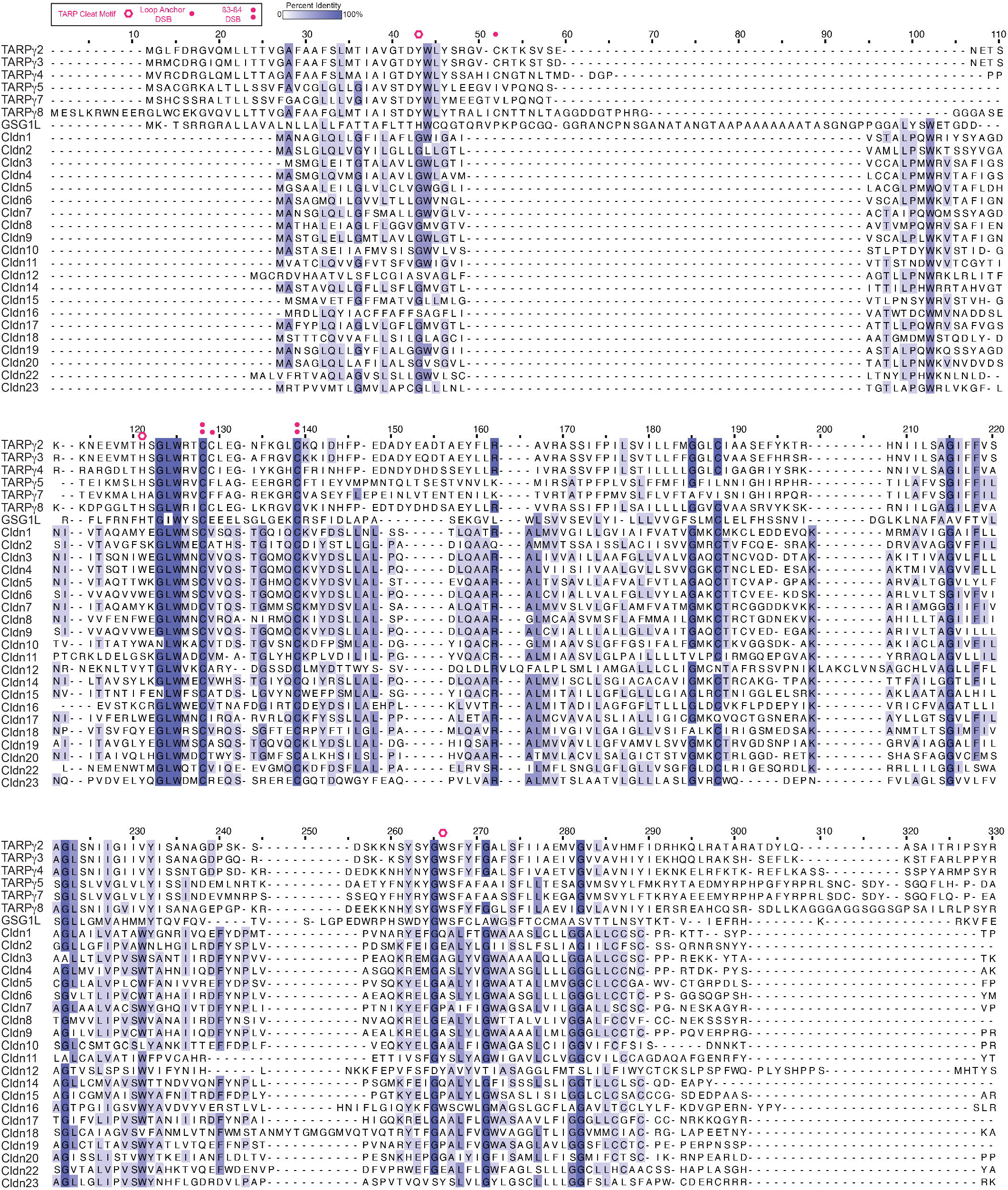
Multiple sequence alignment of TARPs, GSG1L and Claudins. Multiple sequence alignments of TARPs, GSG1L and all members of the Claudin family. The TARPs and GSG1L are distinguished from Claudins by the presence of the TARP cleat motif while the β3-β4 DSB is conserved among both TARPs and Claudins.

**Extended Data Figure 3.**
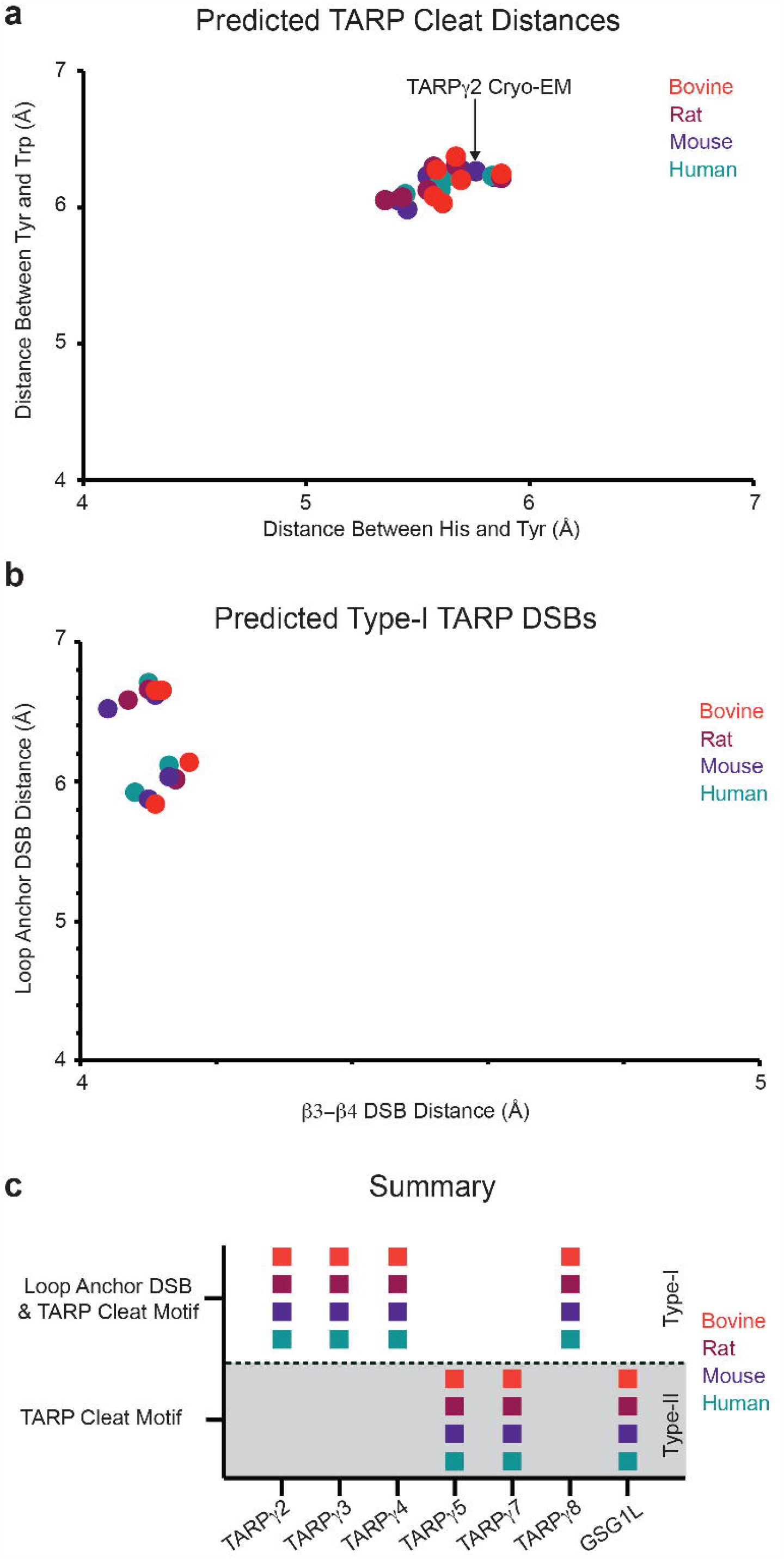
AlphaFold structure prediction of TARPs. a) Conservation of TARP cleat residues in bovine, rat, mouse, and human TARPs. TARPγ2 from this study is pointed out. b) Loop anchor DSB vs. β3-β4 DSB distances. The TARP cleat is predicted to be present in all TARPs. Type-II TARPs are excluded from panel b because the loop anchor DSB is predicted to be absent in type-II TARPs. These findings are summarized in panel c.

